# Nitrogen starvation and stationary phase lipophagy have distinct molecular mechanisms

**DOI:** 10.1101/2020.03.17.996082

**Authors:** Ravinder Kumar, Muhammad Arifur Rahman, Taras Y. Nazarko

## Abstract

In yeast, the selective autophagy of intracellular lipid droplets (LDs) or lipophagy can be induced by either nitrogen (N) starvation or carbon limitation (e.g. in the stationary (S) phase). We developed the yeast, *Komagataella phaffii* (formerly *Pichia pastoris*), as a new lipophagy model and compared the N-starvation and S-phase lipophagy in over 30 autophagy-related mutants using the Erg6-GFP processing assay. Surprisingly, two lipophagy pathways had hardly overlapping stringent molecular requirements. While the N-starvation lipophagy strictly depended on the core autophagic machinery (Atg1-Atg9, Atg18 and Vps15), vacuole fusion machinery (Vam7 and Ypt7) and vacuolar proteolysis (proteinases A and B), only Atg6 and proteinases A and B were essential for the S-phase lipophagy. The rest of the proteins were only partially required in the S-phase. Moreover, we isolated the *prl1* (for positive regulator of lipophagy 1) mutant affected in the S-phase lipophagy but not N-starvation lipophagy. The *prl1* defect was at a stage of delivery of the LDs from the cytoplasm to the vacuole further supporting mechanistically different nature of the two lipophagy pathways. Taken together, our results suggest that N-starvation and S-phase lipophagy have distinct molecular mechanisms.

## Introduction

Autophagy is a highly conserved degradation process in which proteins, protein aggregates and even entire organelles can be sequestered from the cytoplasm by the vacuoles/lysosomes either directly at the vacuolar/lysosomal membrane (microautophagy) or via the double-membrane vesicular intermediates, autophagosomes (macroautophagy) [1],[2]. Autophagy is strongly induced by starvation for nutrients, such as the sources of nitrogen (N), carbon (C) and other elements. The lack of several amino acids can also induce autophagy [3],[4]. Therefore, this process acts as an internal supply of building blocks for cells when the external nutrients become unavailable and allows cells to survive the prolonged periods of starvation.

Lipophagy is an important autophagic process, which delivers the intracellular lipid droplets (LDs) to the vacuoles/lysosomes for degradation and recycling [5]. Lipophagy was initially described in hepatocytes, which become a major site of excessive lipid accumulation in obesity and metabolic syndrome [6]. However, the intracellular lipid metabolism in most eukaryotic cells is also regulated by lipophagy [7], and impaired lipophagy may contribute to the development of many liver and non-liver diseases [8],[9]. Thus, understanding the mechanisms of lipophagy is very important for the prevention and treatment of various lipid accumulation diseases.

The budding yeast *Saccharomyces cerevisiae* was used as a simple lipophagy model by several groups to study the mechanisms of lipophagy. Precisely, lipophagy was induced by either acute N-starvation [10],[11] or C-limitation (either acute [12] or gradual due to the prolonged incubation of cells in the same medium and entering them into stationary (S) phase [13],[11]). These studies described the morphological features of lipophagy and tested the requirements of lipophagy for known autophagy-related (Atg) factors. They suggested that both N-starvation and C-limitation induce microlipophagy [10],[13],[11],[12], the selective microautophagy of LDs, and that this microlipophagy depends on the same core autophagic factors, which are necessary for the formation of autophagic double-membrane in other autophagic pathways [10],[13],[12]. However, such autophagic membrane was never reported to be associated with LDs in the yeast lipophagy studies questioning the role of autophagic machinery in the yeast lipophagy.

Here, we developed the yeast, *Komagataella phaffii* (formerly *Pichia pastoris*), as a new simple model to study lipophagy. The *K. phaffii* has proven to be an excellent model organism for the studies of autophagy-related (Atg) pathways and contributed a lot of mechanistic insights to the field of autophagy [14]. Then, we run the entire collection of *K. phaffii atg*-mutants through the lipophagy assay under both N-starvation and S-phase conditions. As a result, we found that the core autophagic machinery is essential only for the N-starvation lipophagy. The only overlapping stringent molecular requirements for two lipophagy pathways were Atg6 and vacuolar proteinases A and B. In addition, we isolated a new positive regulator of lipophagy 1 (*prl1*) mutant that was deficient only in the S-phase lipophagy. Therefore, we suggest that the N-starvation and S-phase lipophagy pathways have distinct molecular mechanisms.

## Results

### *K. phaffii* is a good model for both N-starvation and S-phase lipophagy

To develop *K. phaffii* as a new lipophagy model, we used the established LD marker protein, Erg6 [10],[12],[11], tagged with the green fluorescent protein (GFP) on the integrative plasmid, pRK2. To confirm the localization of Erg6-GFP to LDs, wild-type (WT) PPY12h cells with pRK2 integrated into the *HIS4* locus were grown in YPD medium for 1 d to an early S-phase and stained with a blue LD dye, monodansylpentane (MDH) [15]. The Erg6-GFP displayed a complete co-localization with MDH (Fig. 1A) suggesting that it is a good LD protein marker for *K. phaffii* under our experimental conditions. The lack of a key Atg protein, Atg8, did not affect the localization of Erg6-GFP to LDs in *atg8* cells (Fig. 1A) making it possible to use the Erg6-GFP as a lipophagy reporter.

**Figure 1.**
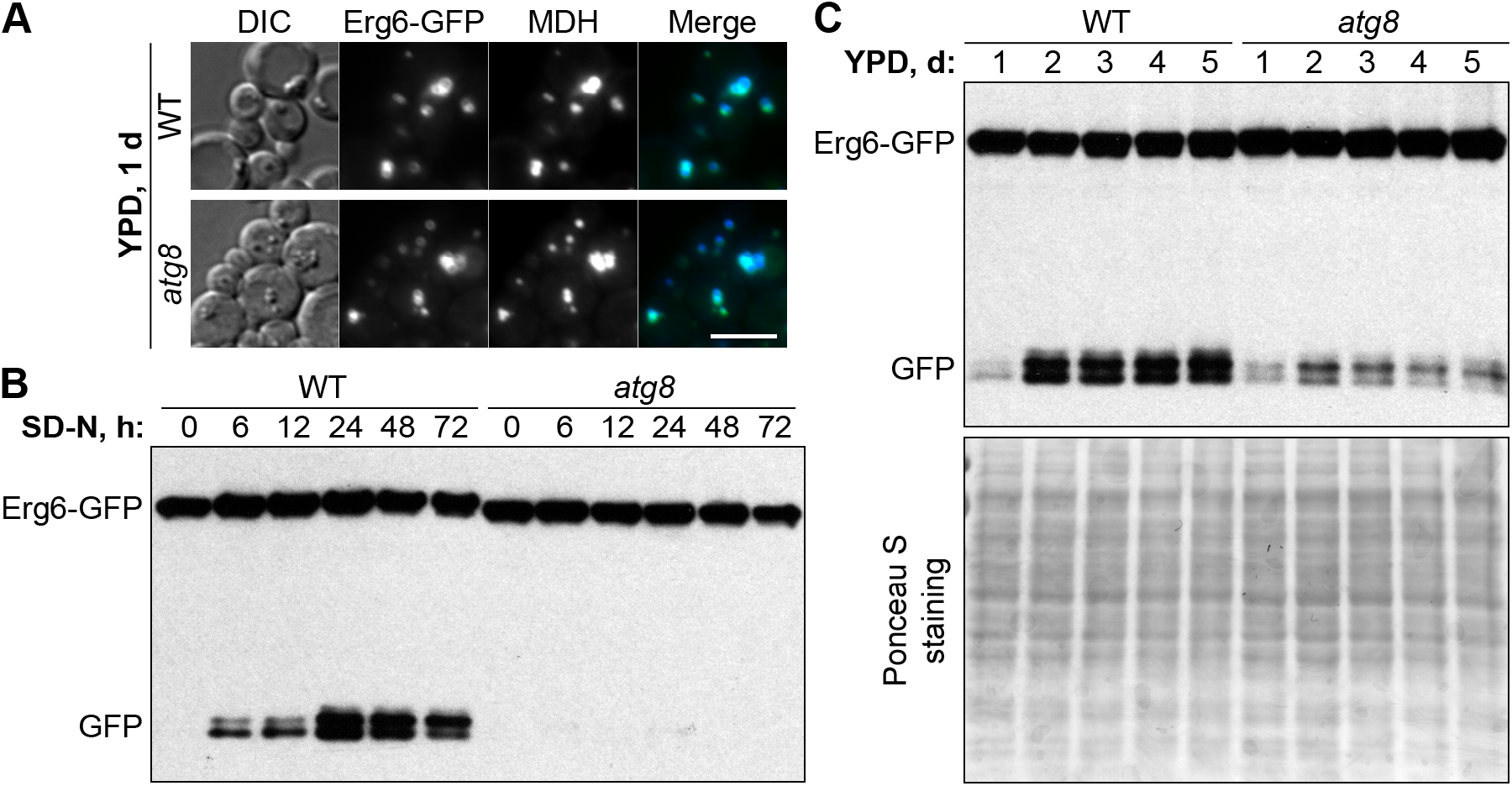
*Komagataella phaffii* is a good model for both N-starvation and S-phase lipophagy. (**A**) *K. phaffii* Erg6-GFP co-localizes with MDH-stained LDs in both WT and *atg8* cells. DIC: differential interference contrast. Scale bar, 5 μm. (**B**) Atg8 is essential for N-starvation lipophagy. Cells were normalized in SD-N at OD_600_ 1 and equal volume of culture (1 mL) was processed at all time-points for both strains to nullify the differential growth (Erg6-GFP dilution) effects in SD-N medium (loading control is not applicable). (**C**) Atg8 is only partially required for S-phase lipophagy. Since biomass slightly decreased during the time-course in S-phase, equal biomass (1 OD_600_) was taken at all time-points for both strains. Ponceau S staining was used as a loading control for S-phase samples.

Then, we developed two Erg6-GFP processing assays to monitor lipophagy: one after the transfer of cells from early S-phase in YPD medium to N-starvation in SD-N medium and another one after the prolonged S-phase in YPD medium. When the LDs with Erg6-GFP are delivered from the cytoplasm to the vacuole, Erg6, but not GFP moiety, is degraded by vacuolar proteases resulting in free GFP, which can be detected by Western blot [16]. The processing of Erg6-GFP to GFP in WT (PPY12h) cells culminated after 24 h of N-starvation (Fig. 1B) and after 2-3 days in YPD medium (Fig. 1C). Therefore, we picked 0 and 24 h, and 1 and 3 d time-points for future N-starvation and S-phase lipophagy assays, respectively.

Interestingly, while *atg8* cells were completely deficient in the Erg6-GFP processing under N-starvation conditions (Fig. 1B), they were only partially compromised in it in S-phase (Fig. 1C) suggesting that N-starvation and S-phase lipophagy pathways might have differences in their molecular requirements. In summary, both N-starvation and S-phase lipophagy pathways were readily induced in *K. phaffii* yeast making it a good model for comparison of their machineries.

### Molecular requirements of N-starvation and S-phase lipophagy in *K. phaffii*

Encouraged by *atg8* results under two lipophagy conditions, we introduced Erg6-GFP into the collection of *K. phaffii* strains deficient in genes that were previously implicated in various Atg-pathways in either *K. phaffii* or other species (Table 2). The collected mutants belong to 4 different WT backgrounds: GS115, GS200, PPY12h and PPY12m. Therefore, we grouped mutants by genetic background and studied them together with the corresponding WT strain, as a control, in both N-starvation and S-phase lipophagy conditions (Figs. 2 and 3, respectively).

**Table 1.**
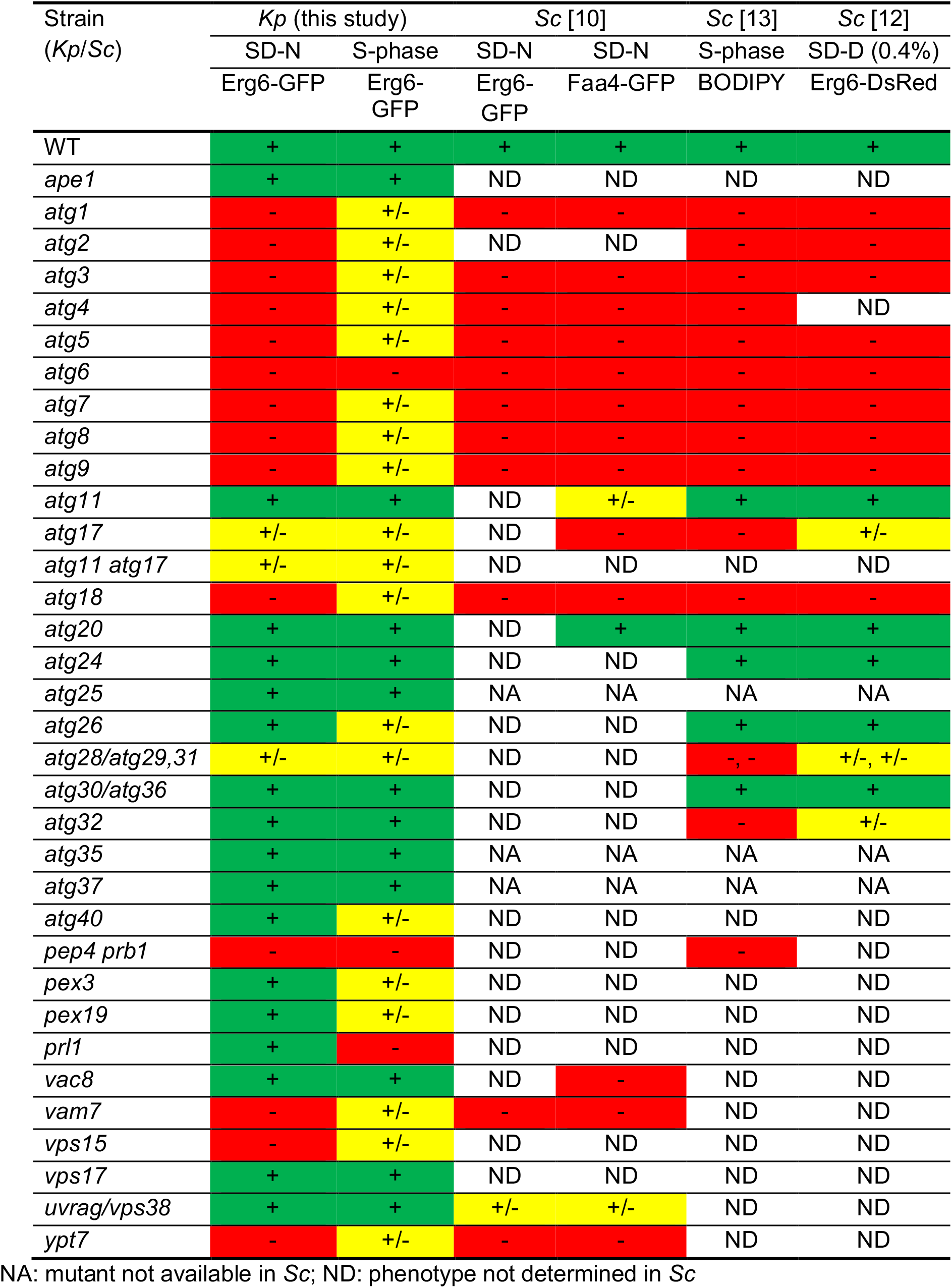
Lipophagy phenotype of *K. phaffii* (this study) and *S. cerevisiae* mutants.

**Table 2.** *K. phaffii* strains used in this study.

| Mutant name | Strain name | Back-ground | Genotype and plasmid | Source |
| --- | --- | --- | --- | --- |
| WT | GS115 | GS115 | *his4* | [22] |
| WT | GS200 | GS200 | *arg4 his4* | [23] |
| WT | PPY12h | PPY12h | *arg4 his4* | [24] |
| WT | PPY12m | PPY12m | *arg4 his4* | [24] |
| *ape1* | SJCF434 | PPY12m | Δ*ape1::Geneticin<sup>R</sup> arg4 his4* | [25] |
| *atg1* | R12 | GS115 | *atg1-1::Zeocin<sup>R</sup> his4* | [26] |
| *atg2* | WDK011 | GS115 | Δ*atg2::Zeocin<sup>R</sup> his4* | [26] |
| *atg3* | *gsa20* | GS115 | *atg3::Zeocin<sup>R</sup> his4* | [26] |
| *atg4* | PPM408 | PPY12h | *atg4::Zeocin<sup>R</sup> arg4 his4* | [27] |
| *atg5* | SJCF2320 | GS115 | Δ*atg5::Zeocin<sup>R</sup> his4* | SL |
| *atg6* | SRDM006 | PPY12m | Δ*atg6::Geneticin<sup>R</sup> arg4 his4* | [28] |
| *atg7* | WDK07 | GS200 | Δ*atg7::ScARG4 arg4 his4* | [29] |
| *atg8* | SJCF925 | PPY12h | Δ*atg8::Geneticin<sup>R</sup> arg4 his4* | [30] |
| *atg9* | R19 | GS115 | *atg9-1::Zeocin<sup>R</sup> his4* | [26] |
| *atg11* | R8 | GS115 | *atg11-2::Zeocin<sup>R</sup> his4* | [31] |
| *atg17* | SJCF929 | PPY12h | Δ*atg17::Geneticin<sup>R</sup> arg4 his4* | [30] |
| *atg11 atg17* | SJCF948 | GS115 | *atg11-2::Zeocin<sup>R</sup> Δatg17::Geneticin<sup>R</sup> his4* | [30] |
| *atg18* | R2 | GS115 | *atg18-1::Zeocin<sup>R</sup> his4* | [26] |
| *atg20* | SRDM020 | PPY12m | Δ*atg20::Geneticin<sup>R</sup> arg4 his4* | SL |
| *atg24* | *paz16* | PPY12h | *atg24::Zeocin<sup>R</sup> arg4 his4* | [32] |
| *atg25* | SJCF1231 | PPY12h | Δ*atg25::Geneticin<sup>R</sup> arg4 his4* | [28] |
| *atg26* | PDG3D | GS200 | Δ*atg26::ScARG4 arg4 his4* | [33] |
| *atg28* | PDG2D | GS200 | Δ*atg28::ScARG4 arg4 his4* | [34] |
| *atg30* | SJCF936 | PPY12h | Δ*atg30::Zeocin<sup>R</sup> arg4 his4* | [30] |
| *atg32* | SJCF1715 | PPY12h | Δ*atg32::Geneticin<sup>R</sup> arg4 his4* | [35] |
| *atg35* | SVN1 | GS200 | Δ*atg35::ScARG4 arg4 his4* | [36] |
| *atg37* | STN96 | PPY12h | Δ*atg37::Geneticin<sup>R</sup> arg4 his4* | [37] |
| *atg40* | SRK2 | PPY12h | Δ*atg40::Zeocin<sup>R</sup> (pRK4) arg4 his4* | This study |
| *pep4 prb1* | SMD1163 | GS115 | *pep4 prb1 his4* | [38] |
| *pex3* | SEW1 | PPY12h | Δ*pex3::PpARG4 arg4 his4* | [39] |
| *pex19* | SKF13 | PPY12h | Δ*pex19::Zeocin<sup>R</sup> arg4 his4* | [40] |
| *prl1* | SRK3 | PPY12h | *prl1::Zeocin<sup>R</sup> (pRK6) arg4 his4* | This study |
| *vac8* | WDY53 | GS200 | Δ*vac8::Zeocin<sup>R</sup> arg4 his4* | [41] |
| *vam7* | SRDM050 | PPY12m | Δ*vam7::Zeocin<sup>R</sup> arg4 his4* | [42] |
| *vps15* | OP5 | GS200 | Δ*vps15::ScARG4 arg4 his4* | [43] |
| *vps17* | SRDM122 | PPY12m | Δ*vps17::Geneticin<sup>R</sup> arg4 his4* | [28] |
| *uvrag* | SRDM083 | PPY12m | Δ*uvrag::Zeocin<sup>R</sup> arg4 his4* | [28] |
| *ypt7* | SRRM197 | PPY12h | Δ*ypt7::Geneticin<sup>R</sup> arg4 his4* | [42] |

SL: Subramani lab

**Figure 2.**
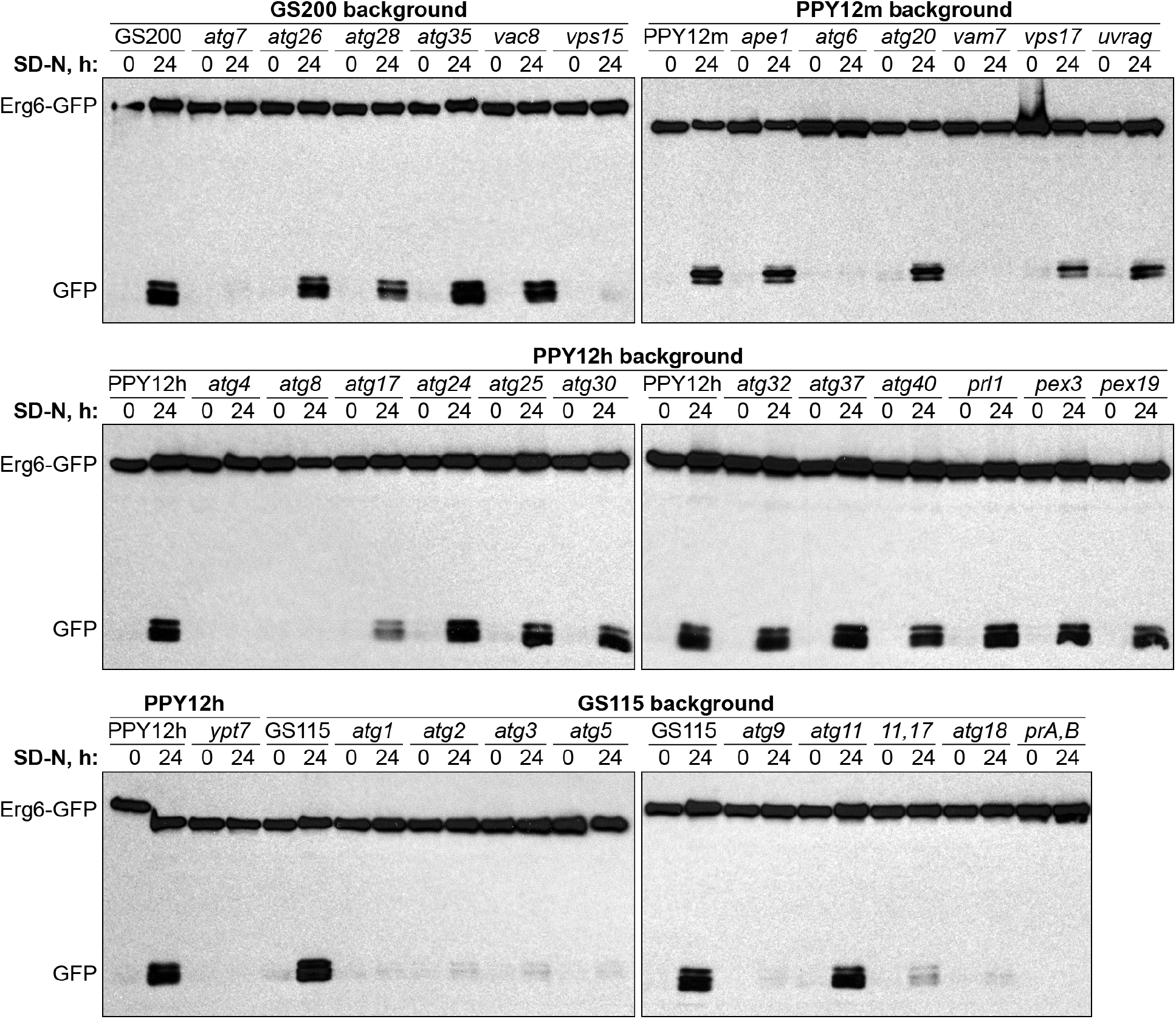
N-starvation lipophagy strictly depends on the core autophagic machinery (Atg1-Atg9, Atg18 and Vps15), vacuole fusion machinery (Vam7 and Ypt7) and vacuolar proteolysis (proteinases A and B). Cells were normalized in SD-N at OD_600_ 1 and equal volume of culture (1 mL) was processed at both time-points for all strains to nullify the differential growth (Erg6-GFP dilution) effects in SD-N medium (loading control is not applicable). *prA,B:* proteinases A and B-deficient mutant *pep4 prb1*.

**Figure 3.**
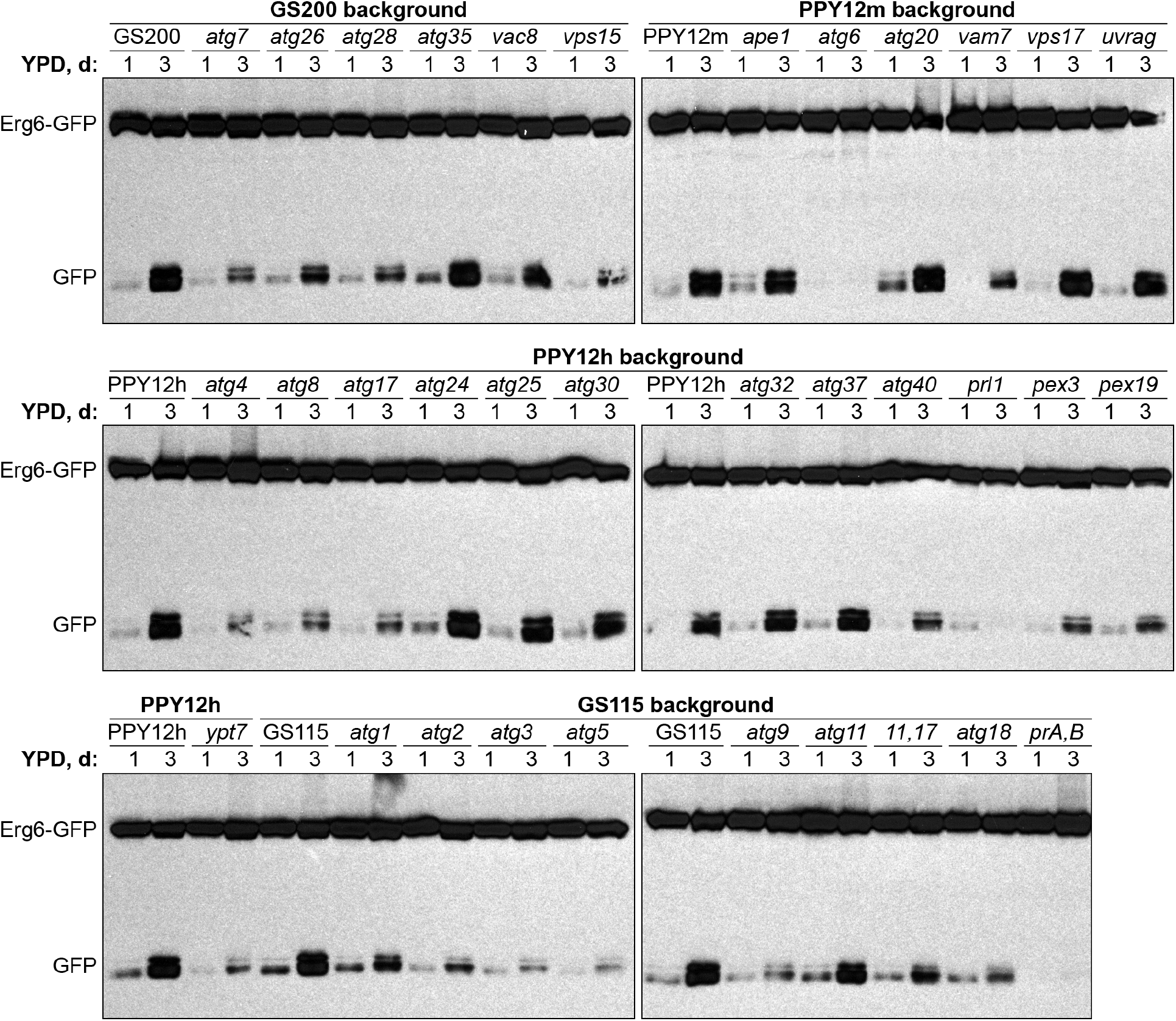
S-phase lipophagy strictly depends only on Atg6, Prl1 (positive regulator of lipophagy 1) and vacuolar proteolysis (proteinases A and B). Since biomass slightly decreased after 3 days in S-phase, equal biomass (1 OD_600_) was taken at both time-points for all strains. Ponceau S staining was used as a loading control and displayed in Fig. S1. *prA,B:* proteinases A and B-deficient mutant *pep4 prb1*.

We found that most of the mutants were either fully deficient (*atg1-atg9*, *atg18*, *pep4 prb1*, *vam7*, *vps15* and *ypt7*) or fully proficient (*ape1*, *atg11*, *atg20*, *atg24-atg26*, *atg30*, *atg32*, *atg35*, *atg37*, *atg40*, *pex3*, *pex19*, *prl1*, *vac8*, *vps17* and *uvrag*) in the Erg6-GFP processing under N-starvation conditions. Only 3 strains (*atg17*, *atg11 atg17* and *atg28*) had an intermediate phenotype (Fig. 2 and Table 1). These results suggested that N-starvation lipophagy strictly depends on the core autophagic machinery (Atg1-Atg9, Atg18 and Vps15), vacuole fusion machinery (Vam7 and Ypt7) and vacuolar proteolysis (proteinases A and B).

In contrast, the Erg6-GFP processing in S-phase was fully deficient in only 3 strains (*atg6*, *pep4 prb1* and *prl1*). It was fully proficient in nearly as many strains (*ape1*, *atg11*, *atg20*, *atg24, atg25*, *atg30*, *atg32*, *atg35*, *atg37*, *vac8*, *vps17* and *uvrag*), as under N-starvation conditions. However, most of the mutants (*atg1-atg5*, *atg7-atg9*, *atg17*, *atg11 atg17*, *atg18*, *atg26*, *atg28*, *atg40*, *pex3*, *pex19*, *vam7*, *vps15* and *ypt7*) had an intermediate phenotype (Figs. 3 and S1; Table 1). Therefore, we concluded that S-phase lipophagy strictly depends only on Atg6, Prl1 and vacuolar proteolysis (proteinases A and B). Summarizing, the N-starvation and S-phase lipophagy pathways have different molecular requirements.

### Prl1 is essential for the delivery of LDs to the vacuole in S-phase

To probe further into the differences between N-starvation and S-phase lipophagy machineries, we took advantage of the *prl1* mutant. This mutant was obtained by the integration of *Zeocin^R^* cassette from the pRK6 plasmid into the genome of PPY12h WT strain (Table 2). The *prl1* mutant displayed a unique phenotype in the screening above: it was fully proficient in the N-starvation lipophagy, but fully deficient in the S-phase lipophagy (Table 1).

To compare the phenotypes of *prl1* cells in the same experiment, we split the cultures of WT, *prl1* and *pep4 prb1* cells after 1 d in YPD medium: small aliquots were transferred to SD-N medium (for 0 and 24 h time-points), while the rest remained in YPD medium (for 1 and 3 d time-points) (Figs. 4A and S2). While N-starvation and S-phase lipophagy pathways were equally well induced in WT cells, they were fully blocked in the proteinases A and B-deficient mutant. Despite *prl1* cells were indistinguishable from WT cells under N-starvation conditions, they were indeed incapable to degrade LDs in S-phase (Fig. 4A). Therefore, we concluded that *prl1* mutant is specifically deficient in the S-phase lipophagy.

**Figure 4.**
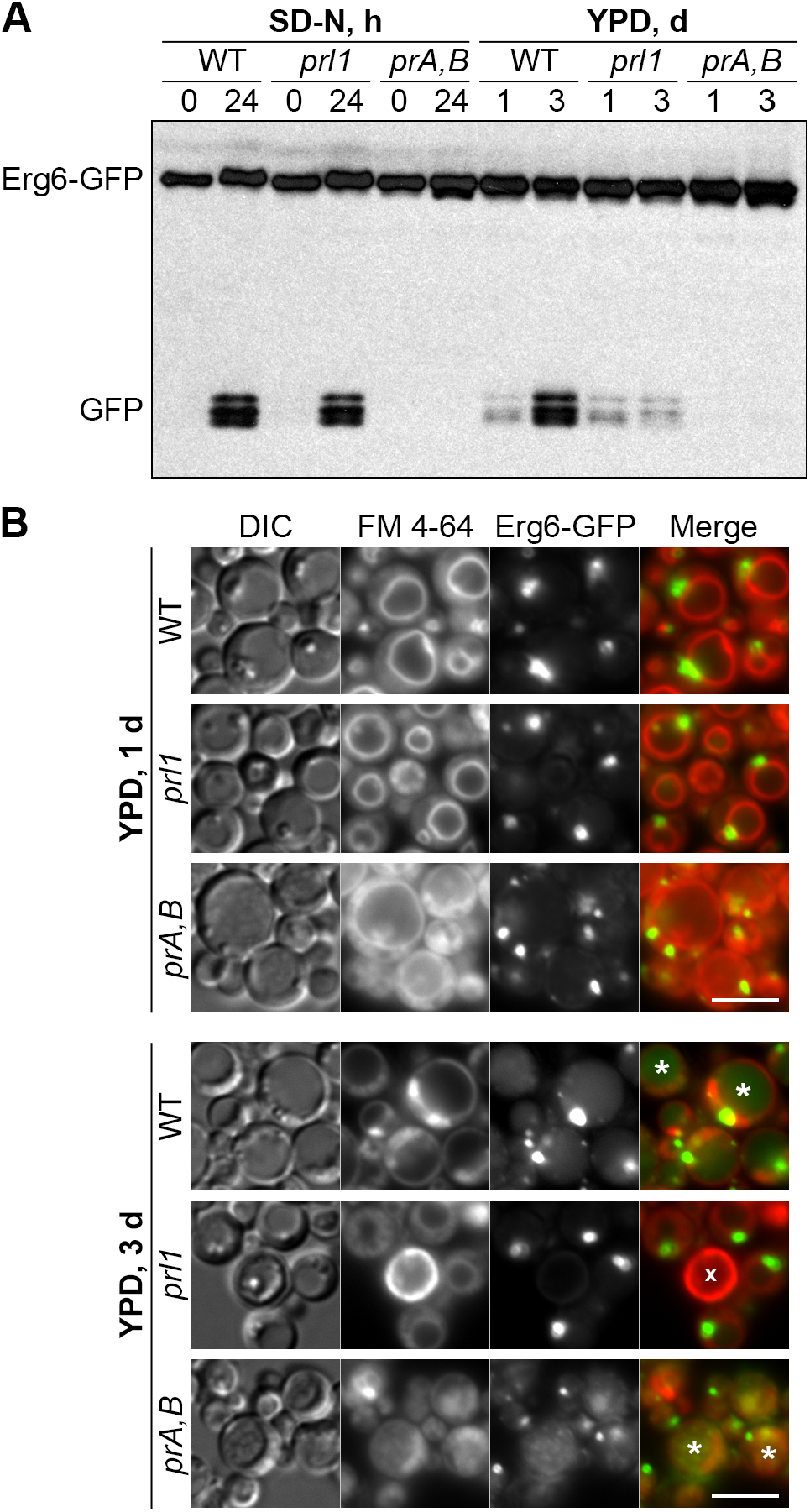
Prl1 is essential for the delivery of LDs to the vacuole in S-phase. (**A**) S-phase but not N-starvation lipophagy depends on Prl1. N-starvation and S-phase cells were processed as described in Figs. 2 and 3, respectively. Ponceau S staining (Fig. S2) was used as a loading control for S-phase samples. (**B**) Prl1 is required for the delivery of LDs to the vacuole in S-phase. Vacuole membranes were stained red with FM 4-64. DIC: differential interference contrast; *prA,B:* proteinases A and B-deficient mutant *pep4 prb1*; *: GFP-containing vacuole; x: dead cell. Scale bar, 5 μm.

To get insight into the step of S-phase lipophagy affected in *prl1* cells, we studied S-phase lipophagy by fluorescence microscopy. The WT, *prl1* and *pep4 prb1* cells with Erg6-GFP reporter were incubated in the YPD medium with FM 4-64, the lipophilic dye that stains specifically the vacuolar membrane. After 1 d, the cells of all strains had LDs outside the vacuole and no GFP fluorescence inside the vacuolar lumen. However, after 3 d, WT and *pep4 prb1* cells developed a diffuse or grainy GFP fluorescence in the vacuolar lumen, respectively (Fig. 4B). Those patterns of luminal GFP fluorescence were consistent with disintegration of the LD-containing autophagic bodies in WT vacuoles and Brownian movement of intact LD-containing autophagic bodies in the proteinases A and B-deficient vacuoles. The *prl1* cells did not gain any GFP fluorescence in the vacuolar lumen after 3 d in YPD suggesting that Prl1 is required for the delivery of LDs to the vacuole in S-phase.

## Discussion

In this study, we introduced *K. phaffii* yeast as a new lipophagy model and compared the lipophagy requirements of *K. phaffii* cells under two conditions: N-starvation and S-phase, the two most popular ways to induce Atg-pathways in yeast. Previous screenings in *S. cerevisiae* done for each of these conditions separately indicated that both of them induced microlipophagy that strongly depended on the core autophagic machinery [10],[13]. However, by comparing the N-starvation and S-phase conditions in the same study with *K. phaffii*, we observed a clear difference in lipophagy requirements (Table 1).

Both the N-starvation and S-phase lipophagy pathways strictly depended on the proteinases A and B consistent with the vacuolar degradation of LDs under the two conditions. While the N-starvation lipophagy strongly relied on the core autophagic machinery represented by Atg1-Atg9, Atg18 and Vps15, the S-phase lipophagy was fully deficient only without the Atg6 protein. Interestingly, only Atg6 and not the other components of the phosphatidylinositol 3-kinase complex I (i.e. Atg14, Atg38, Vps15 and Vps34) stably localized to the vacuolar membrane under both acute and gradual (S-phase) C-limitation conditions in *S. cerevisiae* [12]. Atg6 was necessary for the formation of raft-like domains in the vacuolar membrane [12] that are essential for microlipophagy [13]. Combined, this and previous studies suggest that Atg6 plays a unique role in the S-phase lipophagy, which is different from its established function in the biogenesis of autophagic double-membrane under N-starvation conditions [17].

While a reason for the essential role of the core autophagic machinery in the N-starvation lipophagy is unclear, there is a plausible explanation for the partial requirement of the core autophagic machinery in the S-phase lipophagy. In S-phase, the core autophagic machinery is essential for the correct vacuolar localization of Niemann-Pick type C proteins, Ncr1 and Npc2, that transport sterol from the vacuolar lumen to the vacuolar membrane for the raft-like domain formation [11]. However, the requirements of Ncr1 and Npc2 for (1) raft-like domains formation, (2) their internalization as microautophagic bodies and (3) S-phase lipophagy are partial [11]. Therefore, the core autophagic machinery has, at the end, a partial role in the S-phase lipophagy. Interestingly, it is not required for the correct vacuolar localization of Ncr1 and Npc2 under the N-starvation conditions. Thus, the mechanistic role of the core autophagic machinery in the N-starvation lipophagy is still unknown.

Recently, it was reported that the vacuolar membrane protein, Vph1, which is normally excluded from the raft-like domains in S-phase [18], is also degraded, like LDs, by microautophagy [19]. Interestingly, the S-phase microautophagy of Vph1 was independent of the core autophagic machinery, but relied on the machinery of ESCRT, the endosomal sorting complex required for transport. The same study also reported that the S-phase microautophagy of LDs was partially independent of the core autophagic factor, Atg1, but strongly relied on the ESCRT component, Vps27 [19]. Our results are consistent with these lipophagy observations and extend them to the entire core autophagic machinery being only partially required specifically in the S-phase. However, it is still unclear how LDs and Vph1 can utilize the same microautophagy pathway in S-phase, since they are associated with different vacuolar membrane domains, the raft-like liquid ordered domain and the liquid disordered domain, respectively. Since the S-phase microautophagy of Vph1 does not require Atg6 [19], and the S-phase lipophagy strongly depends on it (Fig. 3), we propose that these pathways have both common (Vps27) and unique (Atg6) requirements.

Our study has also suggested a unique molecular requirement of the S-phase lipophagy versus N-starvation lipophagy, the positive regulator of lipophagy 1 (Prl1). The *prl1* mutant isolated in this study was deficient in lipophagy only in the S-phase. Moreover, we showed that the *prl1’s* lipophagy block is at a trafficking step, since LDs were not delivered from the cytoplasm to the vacuole for degradation in the S-phase. It will be interesting to determine the gene responsible for *prl1* phenotype, since it can help us to further distinguish the molecular mechanisms of these two clearly distinct lipophagy pathways, the N-starvation and S-phase lipophagy.

## Materials and Methods

### Strains and plasmids

The *K. phaffii* strains used in this study are shown in Table 2. These strains were transformed by electroporation [20] with the EcoNI-linearized (R0521S; New England Biolabs) pRK2 plasmid. This plasmid contained the Erg6-GFP expression cassette (for lipophagy studies) and *HIS4* marker gene (for integration into *his4* mutant allele of the recipient strains and selection of His^+^-transformants). The resulting transformants had the following genotype: *his4*∷pRK2 (P_*ERG6*_-*ERG6-GFP*, *HIS4*). They were selected on SD+CSM-His plates (1.7 g/L YNB without amino acids and ammonium sulfate, 20 g/L dextrose, 5 g/L ammonium sulfate, 0.78 g/L CSM-His and 20 g/L agar) and screened for expression of Erg6-GFP by Western blot with anti-GFP bodies (11814460001; Roche) and for localization of Erg6-GFP to LDs by fluorescence microscopy.

### Fluorescence microscopy

Cells were grown for 1 and/or 3 d in culture tubes with 1 mL of YPD medium (10 g/L yeast extract, 20 g/L peptone and 20 g/L dextrose; the autoclaved solution of yeast extract and peptone was mixed with the filter-sterilized 20x solution of dextrose). LDs were stained with 1 μL of 0.1 M MDH solution (SM1000a; Abcepta) during the last 1 h of incubation of cells in YPD medium. Vacuolar membranes were stained with 1 μL of 1 mg/mL solution of FM 4-64 (T3166; Invitrogen) in DMSO added at the beginning of incubation of cells in YPD medium. Then, cells were immobilized on slides using 1% low-melt agarose. For this, the 2 μL drop of cell culture on slide was mixed with the 5 μL drop of 1% low-melt agarose (37°C) on coverslip. Microscopy was done at the Axioskop 2 MOT microscope equipped with the Plan-Apochromat 100x/1.40 NA oil DIC objective and operated by the AxioVision software (all from Carl Zeiss). All the experiments were done at least in duplicate.

### Biochemical studies

Cells were grown in culture tubes with 1 mL of YPD medium and 1 OD_600_ of cells was taken at 1 and 3 d time-points for studies of S-phase lipophagy. For studies of N-starvation lipophagy, 3 OD_600_ of cells were taken at 1 d time-point in YPD medium, washed twice with 1 mL of 1x YNB solution (1.7 g/L YNB without amino acids and ammonium sulfate) and resuspended in 3 mL of SD-N medium (1.7 g/L YNB without amino acids and ammonium sulfate, and 20 g/L dextrose). Then, 1 mL of cell culture was taken at 0 and 24 h time-points in SD-N medium. Both YPD (1 and 3 d) and SD-N (0 and 24 h) samples were TCA precipitated [21] and analyzed by Western blot with anti-GFP bodies (11814460001; Roche). All the experiments were done at least in duplicate.

## Abbreviations

Atg: autophagy-related
C: carbon
DIC: differential interference contrast
ESCRT: endosomal sorting complex required for transport
GFP: green fluorescent protein
LD: lipid droplet
MDH: monodansylpentane
N: nitrogen
Prl1: positive regulator of lipophagy 1
prA,B: proteinases A and B
S: stationary
WT: wild-type

## Acknowledgements

We thank Jean-Claude Farré and Suresh Subramani for strains and helpful discussions.

## Disclosure Statement

No potential conflict of interest was reported by the authors.

## Funding

This work was supported by the NIH grants, DK106344 and GM119571, to TYN.

## References

1. Klionsky DJ. Autophagy: from phenomenology to molecular understanding in less than a decade. Nat Rev Mol Cell Biol. 2007 Nov;8(11):931–7.

2. Ohsumi Y. Historical landmarks of autophagy research. Cell Res. 2014 Jan;24(1):9–23.

3. Takeshige K, Baba M, Tsuboi S, et al. Autophagy in yeast demonstrated with proteinase-deficient mutants and conditions for its induction. J Cell Biol. 1992 Oct;119(2):301–11.

4. Sutter BM, Wu X, Laxman S, et al. Methionine inhibits autophagy and promotes growth by inducing the SAM-responsive methylation of PP2A. Cell. 2013 Jul 18;154(2):403–15.

5. Singh R, Kaushik S, Wang Y, et al. Autophagy regulates lipid metabolism. Nature. 2009 Apr 30;458(7242):1131–5.

6. Marchesini G, Brizi M, Bianchi G, et al. Nonalcoholic fatty liver disease: a feature of the metabolic syndrome. Diabetes. 2001 Aug;50(8):1844–50.

7. Singh R, Cuervo AM. Autophagy in the cellular energetic balance. Cell Metab. 2011 May 4;13(5):495–504.

8. Zhang Z, Yao Z, Chen Y, et al. Lipophagy and liver disease: New perspectives to better understanding and therapy. Biomed Pharmacother. 2018 Jan;97:339–348.

9. Zhou K, Yao P, He J, et al. Lipophagy in nonliver tissues and some related diseases: Pathogenic and therapeutic implications. J Cell Physiol. 2019 Jun;234(6):7938–7947.

10. van Zutphen T, Todde V, de Boer R, et al. Lipid droplet autophagy in the yeast Saccharomyces cerevisiae. Mol Biol Cell. 2014 Jan;25(2):290–301.

11. Tsuji T, Fujimoto M, Tatematsu T, et al. Niemann-Pick type C proteins promote microautophagy by expanding raft-like membrane domains in the yeast vacuole. Elife. 2017 Jun 7;6.

12. Seo AY, Lau PW, Feliciano D, et al. AMPK and vacuole-associated Atg14p orchestrate mu-lipophagy for energy production and long-term survival under glucose starvation. Elife. 2017 Apr 10;6.

13. Wang CW, Miao YH, Chang YS. A sterol-enriched vacuolar microdomain mediates stationary phase lipophagy in budding yeast. J Cell Biol. 2014 Aug 4;206(3):357–66.

14. Farre JC, Subramani S. Mechanistic insights into selective autophagy pathways: lessons from yeast. Nat Rev Mol Cell Biol. 2016 Sep;17(9):537–52.

15. Yang HJ, Hsu CL, Yang JY, et al. Monodansylpentane as a blue-fluorescent lipid-droplet marker for multi-color live-cell imaging. PLoS One. 2012;7(3):e32693.

16. Shintani T, Klionsky DJ. Cargo proteins facilitate the formation of transport vesicles in the cytoplasm to vacuole targeting pathway. J Biol Chem. 2004 Jul 16;279(29):29889–94.

17. Obara K, Ohsumi Y. PtdIns 3-Kinase Orchestrates Autophagosome Formation in Yeast. J Lipids. 2011;2011:498768.

18. Toulmay A, Prinz WA. Direct imaging reveals stable, micrometer-scale lipid domains that segregate proteins in live cells. J Cell Biol. 2013 Jul 8;202(1):35–44.

19. Oku M, Maeda Y, Kagohashi Y, et al. Evidence for ESCRT- and clathrin-dependent microautophagy. J Cell Biol. 2017 Oct 2;216(10):3263–3274.

20. Cregg JM, Russell KA. Transformation. Methods Mol Biol. 1998;103:27–39.

21. Baerends RJ, Faber KN, Kram AM, et al. A stretch of positively charged amino acids at the N terminus of Hansenula polymorpha Pex3p is involved in incorporation of the protein into the peroxisomal membrane. J Biol Chem. 2000 Apr 7;275(14):9986–95.

22. Cregg JM, Barringer KJ, Hessler AY, et al. Pichia pastoris as a host system for transformations. Mol Cell Biol. 1985 Dec;5(12):3376–85.

23. Waterham HR, de Vries Y, Russel KA, et al. The Pichia pastoris PER6 gene product is a peroxisomal integral membrane protein essential for peroxisome biogenesis and has sequence similarity to the Zellweger syndrome protein PAF-1. Mol Cell Biol. 1996 May;16(5):2527–36.

24. Gould SJ, McCollum D, Spong AP, et al. Development of the yeast Pichia pastoris as a model organism for a genetic and molecular analysis of peroxisome assembly. Yeast. 1992 Aug;8(8):613–28.

25. Farre JC, Vidal J, Subramani S. A cytoplasm to vacuole targeting pathway in P. pastoris. Autophagy. 2007 May-Jun;3(3):230–4.

26. Stromhaug PE, Bevan A, Dunn WA, Jr. GSA11 encodes a unique 208-kDa protein required for pexophagy and autophagy in Pichia pastoris. J Biol Chem. 2001 Nov 9;276(45):42422–35.

27. Mukaiyama H, Oku M, Baba M, et al. Paz2 and 13 other PAZ gene products regulate vacuolar engulfment of peroxisomes during micropexophagy. Genes Cells. 2002 Jan;7(1):75–90.

28. Farre JC, Mathewson RD, Manjithaya R, et al. Roles of Pichia pastoris Uvrag in vacuolar protein sorting and the phosphatidylinositol 3-kinase complex in phagophore elongation in autophagy pathways. Autophagy. 2010 Jan;6(1):86–99.

29. Yuan W, Stromhaug PE, Dunn WA, Jr. Glucose-induced autophagy of peroxisomes in Pichia pastoris requires a unique E1-like protein. Mol Biol Cell. 1999 May;10(5):1353–66.

30. Nazarko TY, Farre JC, Subramani S. Peroxisome size provides insights into the function of autophagy-related proteins. Mol Biol Cell. 2009 Sep;20(17):3828–39.

31. Kim J, Kamada Y, Stromhaug PE, et al. Cvt9/Gsa9 functions in sequestering selective cytosolic cargo destined for the vacuole. J Cell Biol. 2001 Apr 16;153(2):381–96.

32. Ano Y, Hattori T, Oku M, et al. A sorting nexin PpAtg24 regulates vacuolar membrane dynamics during pexophagy via binding to phosphatidylinositol-3-phosphate. Mol Biol Cell. 2005 Feb;16(2):446–57.

33. Stasyk OV, Nazarko TY, Stasyk OG, et al. Sterol glucosyltransferases have different functional roles in Pichia pastoris and Yarrowia lipolytica. Cell Biol Int. 2003;27(11):947–52.

34. Stasyk OV, Stasyk OG, Mathewson RD, et al. Atg28, a novel coiled-coil protein involved in autophagic degradation of peroxisomes in the methylotrophic yeast Pichia pastoris. Autophagy. 2006 Jan-Mar;2(1):30–8.

35. Farre JC, Burkenroad A, Burnett SF, et al. Phosphorylation of mitophagy and pexophagy receptors coordinates their interaction with Atg8 and Atg11. EMBO Rep. 2013 May;14(5):441–9.

36. Nazarko VY, Nazarko TY, Farre JC, et al. Atg35, a micropexophagy-specific protein that regulates micropexophagic apparatus formation in Pichia pastoris. Autophagy. 2011 Apr;7(4):375–85.

37. Nazarko TY, Ozeki K, Till A, et al. Peroxisomal Atg37 binds Atg30 or palmitoyl-CoA to regulate phagophore formation during pexophagy. J Cell Biol. 2014 Feb 17;204(4):541–57.

38. Tuttle DL, Dunn WA, Jr. Divergent modes of autophagy in the methylotrophic yeast Pichia pastoris. J Cell Sci. 1995 Jan;108 (Pt 1):25–35.

39. Wiemer EA, Luers GH, Faber KN, et al. Isolation and characterization of Pas2p, a peroxisomal membrane protein essential for peroxisome biogenesis in the methylotrophic yeast Pichia pastoris. J Biol Chem. 1996 Aug 2;271(31):18973–80.

40. Snyder WB, Faber KN, Wenzel TJ, et al. Pex19p interacts with Pex3p and Pex10p and is essential for peroxisome biogenesis in Pichia pastoris. Mol Biol Cell. 1999 Jun;10(6):1745–61.

41. Chang T, Schroder LA, Thomson JM, et al. PpATG9 encodes a novel membrane protein that traffics to vacuolar membranes, which sequester peroxisomes during pexophagy in Pichia pastoris. Mol Biol Cell. 2005 Oct;16(10):4941–53.

42. Manjithaya R, Anjard C, Loomis WF, et al. Unconventional secretion of Pichia pastoris Acb1 is dependent on GRASP protein, peroxisomal functions, and autophagosome formation. J Cell Biol. 2010 Feb 22;188(4):537–46.

43. Stasyk OV, van der Klei IJ, Bellu AR, et al. A Pichia pastoris VPS15 homologue is required in selective peroxisome autophagy. Curr Genet. 1999 Nov;36(5):262–9.

