## Supplement for "Nitrogen starvation and stationary phase lipophagy have distinct molecular mechanisms"

**Figure S1**

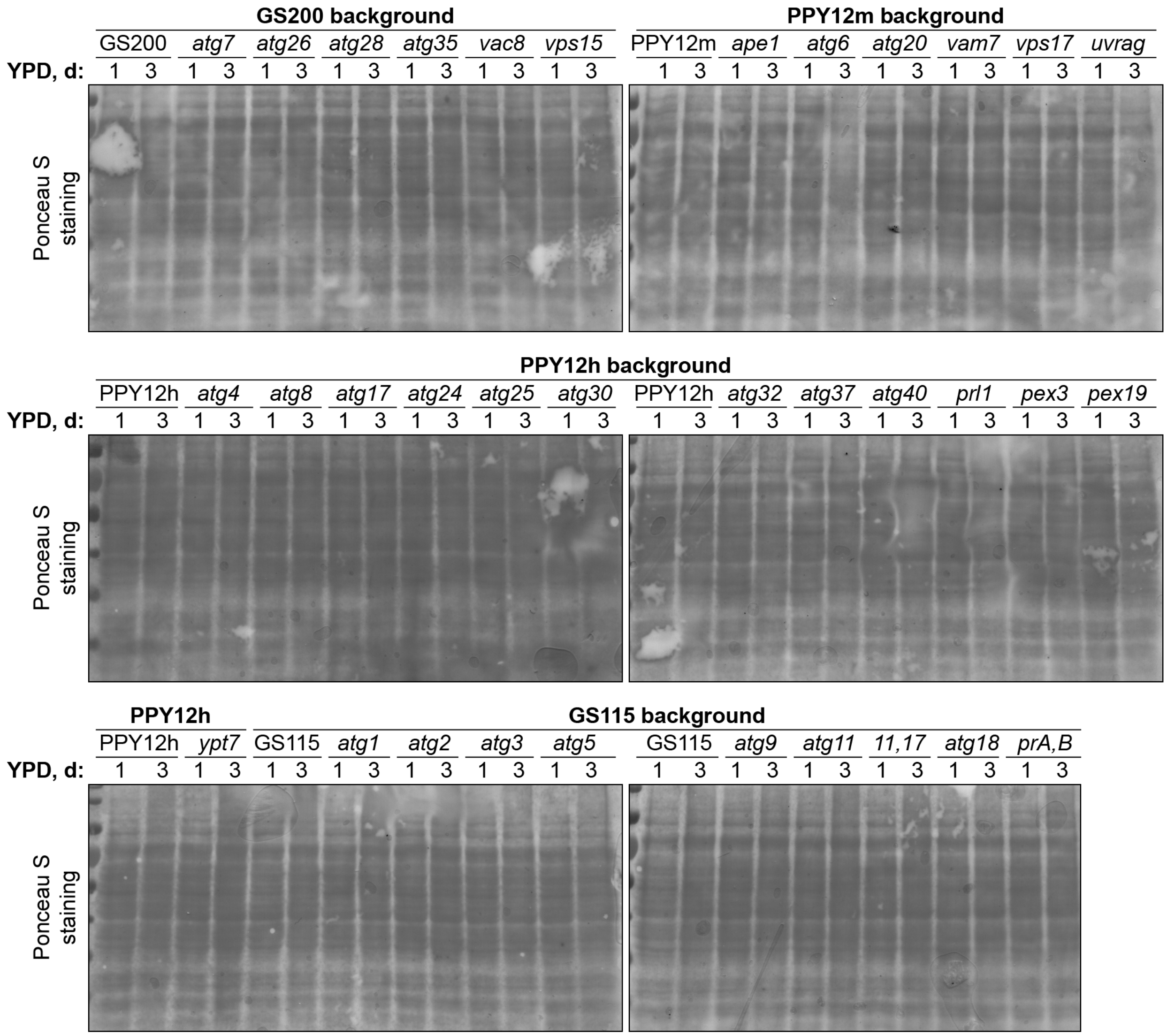


**Figure S1.** Supplemental figure for Fig. 3. Loading control (see Fig. 3 for details).

**Figure S2**

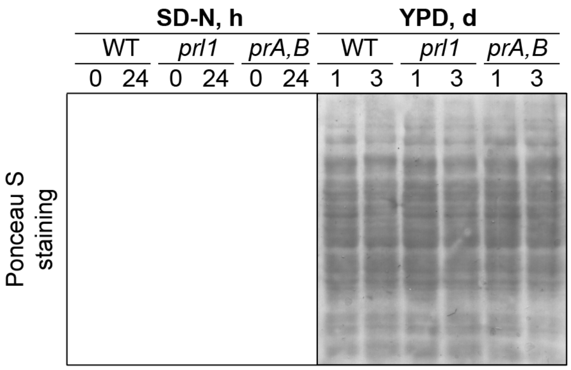


**Figure S2.** Supplemental figure for Fig. 4A. Loading control (see Fig. 4A for details).
